# Metagenomics Study of Bacterial Communities from Yamuna River water of city of Taj, Agra

**DOI:** 10.1101/2022.03.21.485165

**Authors:** Nupur Raghav, Pooja Saraswat, Janendra Nath Srivastava, Rajiv Ranjan

**Affiliations:** Department of Botany, Dayalbagh Educational Institute (Deemed University), Dayalbagh, Agra, India

**Keywords:** Yamuna River, Physico-chemical parameters, APHA, Heavy metals, Metagenomics

## Abstract

Yamuna river water of Agra city is heavily contaminated with toxic pollutants including heavy metal that causes severe damage to ecological and social aspects. At present, the direct use of river water for the purpose of drinking causes severe hazards due to anthropogenic activities causing environmental pollution in rivers. In present study, Yamuna River water was collected from three different sites of Agra city. Various physico-chemical parameters were estimated by following the standard methods of APHA and the concentration of heavy metals were measured using Atomic Absorption Spectrophotometer (AAS). In case of physico-chemical parameters characterization, the obtained values were extremely above the permissible limits. On account of the research work; the Yamuna river water samples contains heavy metal concentrations (Cadmium, Chromium, Copper, Lead and Nickel) above the desirable and admissible levels except for Zinc. During analysis of non-culturable bacteria a substantial bacterial diversity was observed in the Yamuna river water samples collected from different sites. The water samples were subjected to metagenomic analysis using Illumina platform which revealed that Proteobacteria (phylum), Betaproteobacteria (class), Burkholderiales (order), Comamonadaceae (family), *Hydrogenophaga* (genus) and *Chloroflexi bacterium OLB 14* (species) were found as the most dominant bacterial taxonomic abundance in the river water samples. The presence of such bacterial communities in water indicates the availability of pollutants and suggests the futuristic use in the field of bioremediation.

## INTRODUCTION

The quality and quantity of water is gaining extensive attention throughout the world due to massive population growth and increasing trends of social economic development. Water (Blue Gold) one of the most treasured natural resources is responsible for life on earth by contribution, development and growth of human civilization. Most of the civilization all around the world evolved on the rivers banks. Unplanned urbanization, indiscriminate industrialization, rapidly increasing population along the rivers and their catchment area have caused tremendous loss and stress on water quality and their resources.

River Yamuna is the major tributary to River Ganga (India’s largest river) and one of the major rivers in India. Both the rivers cater the fundamental needs of mankind in northern state of India. The extreme cause of pollution in rivers are excessive discharge of domestic waste water from adjacent towns and residents contributing about two-third load of pollution and the rest one third is caused by agricultural and industrial effluents (CPCB 1978). At present, the direct use of river water for the purpose of drinking causes severe hazards due to anthropogenic activities causing environmental pollution in rivers (Barik and Patel 2004).

The deterioration of Yamuna river water of city of Taj, Agra, Uttar Pradesh, India has been a cause of serious concern. Assessment of bacterial communities in the Yamuna River water samples is necessary to ensure the quality of water. Previous techniques of characterization and identification of bacterial isolates was through serial dilution culture based technique and Sanger sequencing. However for the modern researchers, these methods of isolation have lost the charm due to certain limitations i.e. lower sequencing depth and high proportion of non-culturable bacteria (Sundberg et al. 2013; Zhou et al. 2015). It has been proved that metagenomics is a cost effective and time effective identification method that employs the fusion of total genomics DNA isolation from a consortium of species and next generation sequencing (Garrido-Cardenas and Manzano-Agugliaro 2017).

Hence in this study, the attempt was made to analyse the physico-chemical parameters of Yamuna river water and to evaluate the non-culturable bacteria from river water samples through metagenomics approach.

## MATERIAL METHOD

### Study Area

Agra (city of Taj) is located in western U.P. between 27.11’ degree Latitude North and 78.0’ degree to 78.2’ degree Longitude east and its Altitude is 169 meters above sea level (Verma and Singh 2016). The population of Agra city according to the Census (2011) is 1,574,542. The density of city is 11,167 persons per sq. km which is very much high in comparison to state average of 819 persons per sq. km. The total slum population of Agra is 533,554 which are around 33% of the overall population (Census of India 2011). The floating population is recorded as 0.3 million (AJS 2015). As per urbanization level, Agra district is at 8^th^ position among other districts in Uttar Pradesh. The city Agra is located on the bank of river Yamuna.

Yamuna river is one of the largest tributary of the holy Ganga river and one of the most prominent and important rivers of India. It is investigated that around 60.3 million people depend on the river water for their daily needs of life (CPCB 2006). Excessive pollution load due to domestic and industrial wastewater falling from various drains into the Yamuna River reveals that it is continuously increasing day-by-day.

### Site Characterization and Sample Collection

Yamuna river water samples were collected from different sites of Agra city, based upon the existence of industries and disposal of raw sewage which are responsible for main source of hazardous pollutant contamination into the river. The sampling sites are as follows:-S_1_-Kailash Ghat, S_2_- Poiyah Ghat and S_3_-Hathi Ghat (Fig. 1). The Morphometric details along with the description of each site are given in Table 1.

**Table 1.** Morphometric Details of Sampling Sites.

| Sampling Sites | Description | Latitude | Longitude | Elevation |
| --- | --- | --- | --- | --- |
| Kailash Ghat (S<sub>1</sub>) | It is situated about 13 km from the university campus, has steep sloping topography and a temple of Lord Shiva on the River Bank. This site is the upstream of Agra. | 27°14'15.60" N | 77°56'08.02" E | 163 m |
| Poiyah Ghat (S<sub>2</sub>) | It is situated about 5 km from the university campus, has large flat river Bank. Near river bank a cremation ground is located where dead bodies are brought to be burnt. | 27°25'53.01" N | 78°02'18.99" E | 406 m |
| Hathi Ghat (S<sub>3</sub>) | It is situated about 7 km from the university campus. This site is famous for idol immersion. | 27°18'53.65" N | 78°02'57.95" E | 423 m |

**Fig. 1.**
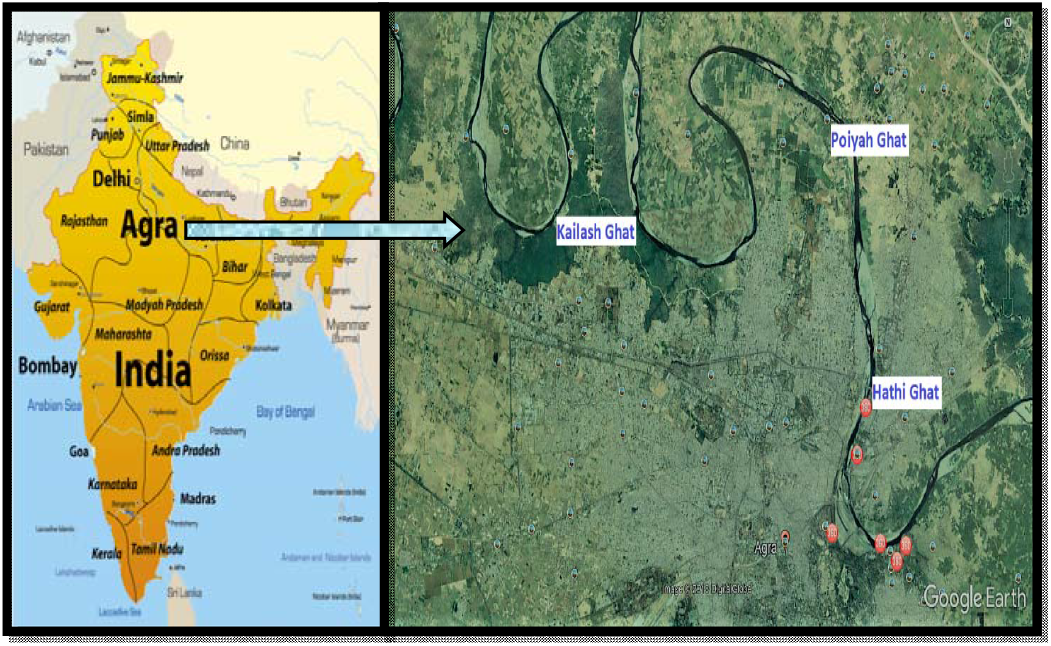
Map showing sampling sites

Water samples from River were collected from three sites Kailash Ghat (N 27°14’15.60” E 77°56’08.02”), Poiyah Ghat (N 27°25’53.01”E 78°02’18.99”) and Hathi Ghat (N 27°18’53.65”E 78°02’57.95”) between 5 to 7 A.M. River water samples were collected in pre-sterilized bottles (5 L capacity) from River banks similar methodology as Behera et al. (2020). Thereafter all the samples were brought to the laboratory for analysis of physicochemical parameters and rest of the samples were stored at 4°C for further analysis and later stored in −80°C for metagenomic analysis.

### Measurement of physico-chemical parameters and heavy metals

The physico-chemical parameters of collected Yamuna river water were measured by following the methodology given by Ambasht (1990) and APHA (1998). To analyse the physical parameters of our water samples, the samples were immediately taken to the laboratory and tested within 2-3 hrs after collection. The pH of samples was taken using pH meter (Ri-Digital-Model-152-R) and dissolved oxygen (mg/L) using titration assembly. Also, the brief details of other parameters and the equipments used in the study are presented in Table 2. All the parameters including alkalinity, hardness, chlorides, dissolved oxygen (DO), BOD, COD and heavy metals were measured in mg/L.

**Table 2.**
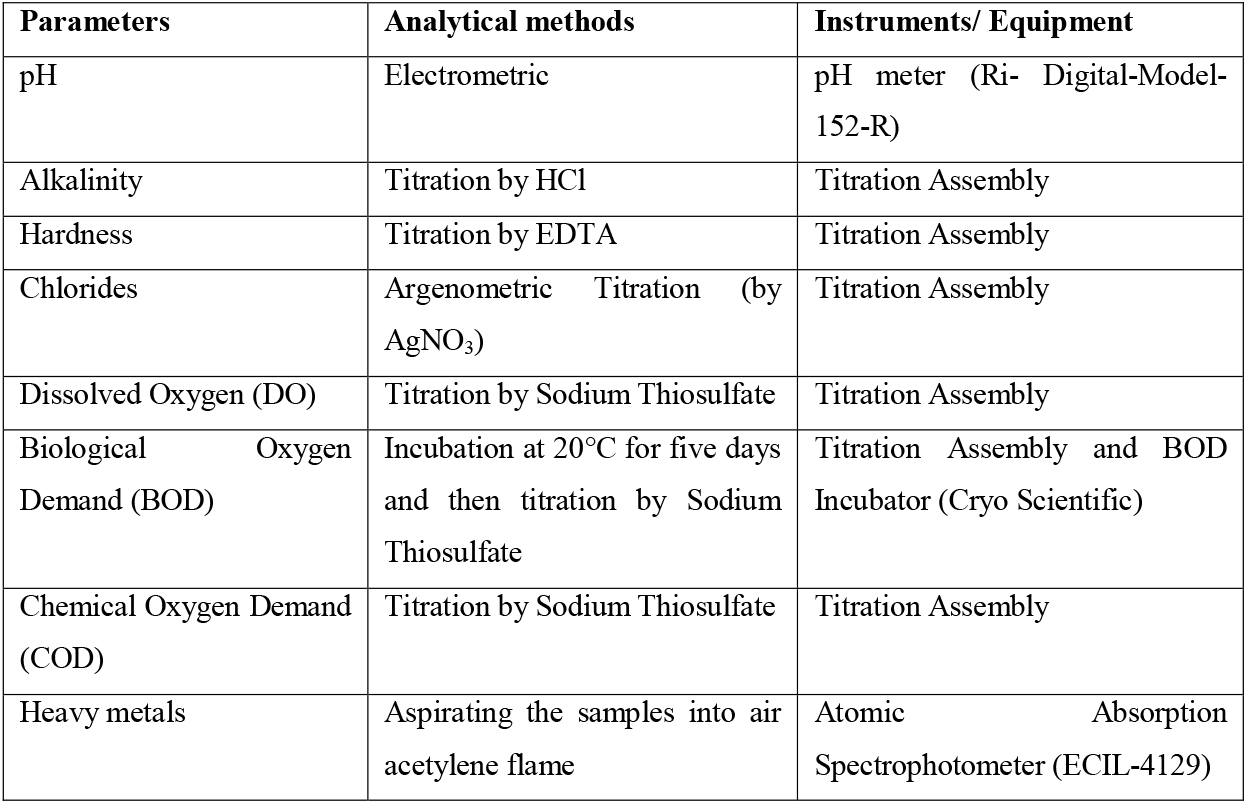
Analytical methods and instruments used in estimating the physico-chemical parameters of Yamuna river water.

## ISOLATION AND IDENTIFICATION OF NON-CULTURABLE BACTERIA

### DNA isolation and estimation

The water samples were sent to Xcelris Labs Ltd. Ahemadabad and DNA was isolated from the sample. Quality of genomic DNA was checked on 0.8% agarose gel (loaded 3μl) for the single intact band. The gel was run at 110 V for 30 mins. Concentration was determined using Qubit® 2.0 Fluorometer through 1 μl of sample.

### Preparation of library

The paired-end sequencing library was prepared using *Truseq Nano DNA Library prep kit*. The library preparation process was initiated with 200 ng g-DNA. The g-DNA was mechanically sheared into smaller fragments by covaris followed by continuous step of end-repair where an ‘A’ is added to the 3’ ends making the DNA fragments ready for adapter ligation. After this stage, platform-specific adapters were ligated to both ends of the DNA fragments so that chimera could generate at low level. The adapters consists sequences that were necessary for binding the dual-barcoded libraries to a flow cell for sequencing purposes, allowing for PCR amplification of adapter-ligated fragments, and binding of standard Illumina sequencing primers. To ensure maximum yields from limited amounts of starting material, a high-fidelity amplification step was performed using HiFi PCR Master Mix. and then the obtained library will be analysed using Bioanalyzer 2100 (Agilent Technologies) with the help of highly sensitive DNA chip. Qubit concentration were obtained for the library and the mean peak size from Bioanalyzer profile, library was loaded onto Illumina platform for cluster generation and sequencing. The library is loaded onto NextSeq to generate cluster and sequencing is done after checking the quality of library produced. The template fragments were allowed by paired-end sequencing to be sequenced in both the directions (forward and reverse). As Paired-end sequencing allows the sequencing of template strand in both directions (forward and reverse) on NextSeq platform.

The library molecules were bind to complementary adapter oligos on paired-end flow cell. Selective cleavage of the forward strands after re-synthesis of the reverse strand during sequencing was allowed by designed adapters. Sequencing from the opposite end of the fragment was done through copied reverse strand.

### Bioinformatics analysis

The next-gen sequencing (NGS) of our sample was done on the Illumina platform using *TruSeq Nano DNA library prep kit*. The read length 2 ×150 PE and < 3GB data/sample. For further metagenomic analysis in the study all the reads were taken to study the taxonomic and functional gene characterisation.

## RESULTS

### Sample collection and physicochemical parameter analysis

Samples were collected from three different sites namely Kailash ghat, Poiyah Ghat and Hathi Ghat for Yamuna river water analysis. For each site samples were collected in three different seasons (Monsoon, winter and summer). The collected samples were tested for the physicochemical parameters which are determined in Table 3. It also represents the average value alkalinity, chlorides, total hardness, dissolved oxygen, Biological Oxygen Demand (BOD), Chemical Oxygen Demand (COD) and heavy metals seasonally.

**Table 3:**
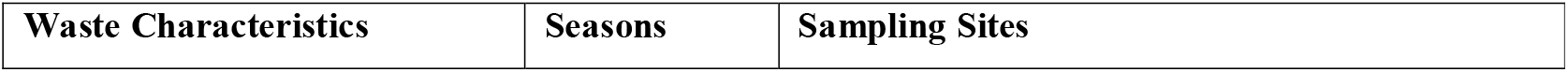
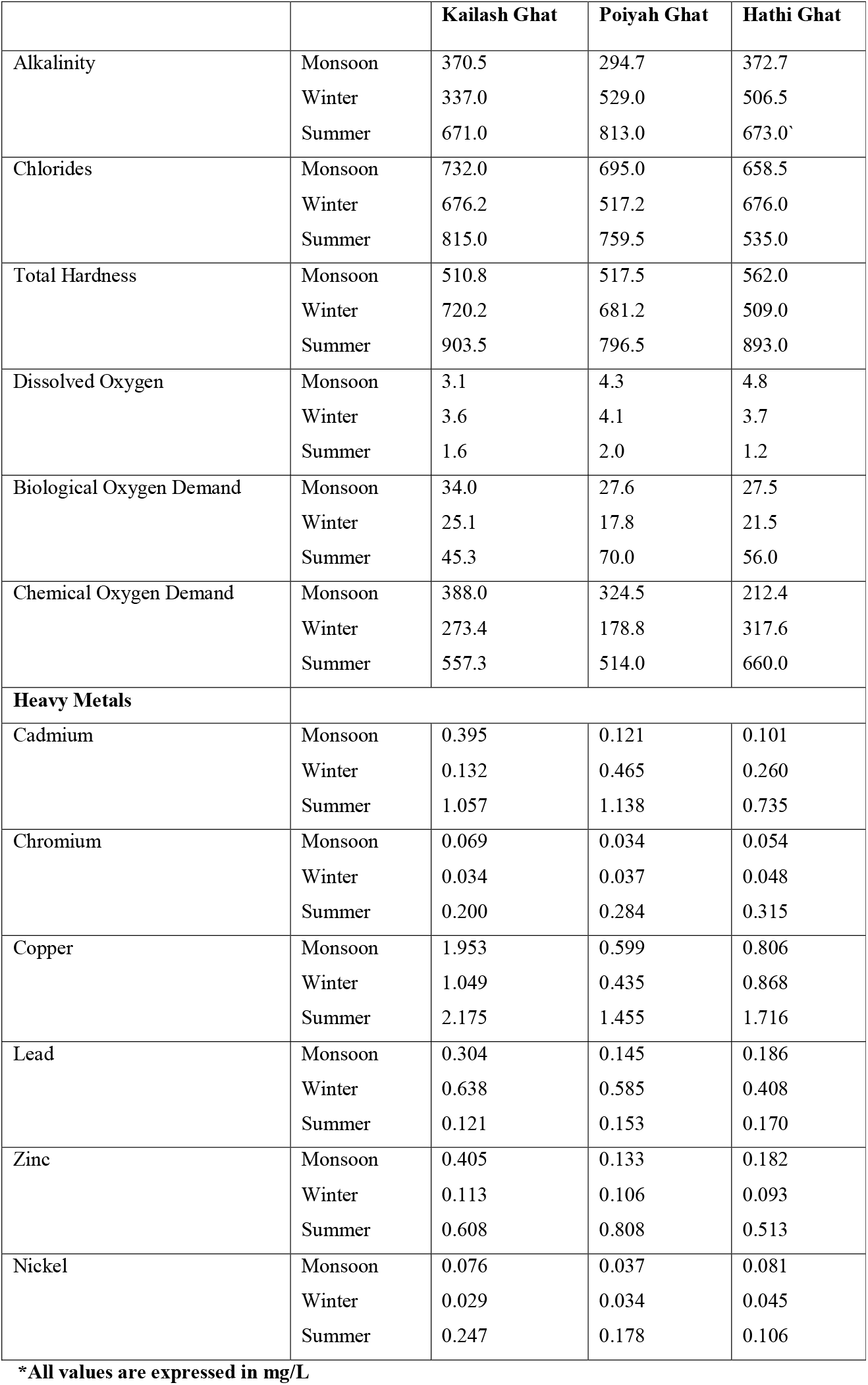
Physico-chemical and Heavy Metals analysis of Yamuna River water samples.

| Waste Characteristics | Seasons | Sampling Sites |
| --- | --- | --- |

|  |  | **Kailash Ghat** | **Poiyah Ghat** | **Hathi Ghat** |
| --- | --- | --- | --- | --- |
| Alkalinity | Monsoon | 370.5 | 294.7 | 372.7 |
|  | Winter | 337.0 | 529.0 | 506.5 |
|  | Summer | 671.0 | 813.0 | 673.0` |
| Chlorides | Monsoon | 732.0 | 695.0 | 658.5 |
|  | Winter | 676.2 | 517.2 | 676.0 |
|  | Summer | 815.0 | 759.5 | 535.0 |
| Total Hardness | Monsoon | 510.8 | 517.5 | 562.0 |
|  | Winter | 720.2 | 681.2 | 509.0 |
|  | Summer | 903.5 | 796.5 | 893.0 |
| Dissolved Oxygen | Monsoon | 3.1 | 4.3 | 4.8 |
|  | Winter | 3.6 | 4.1 | 3.7 |
|  | Summer | 1.6 | 2.0 | 1.2 |
| Biological Oxygen Demand | Monsoon | 34.0 | 27.6 | 27.5 |
|  | Winter | 25.1 | 17.8 | 21.5 |
|  | Summer | 45.3 | 70.0 | 56.0 |
| Chemical Oxygen Demand | Monsoon | 388.0 | 324.5 | 212.4 |
|  | Winter | 273.4 | 178.8 | 317.6 |
|  | Summer | 557.3 | 514.0 | 660.0 |
| **Heavy Metals** |  |  |  |  |
| Cadmium | Monsoon | 0.395 | 0.121 | 0.101 |
|  | Winter | 0.132 | 0.465 | 0.260 |
|  | Summer | 1.057 | 1.138 | 0.735 |
| Chromium | Monsoon | 0.069 | 0.034 | 0.054 |
|  | Winter | 0.034 | 0.037 | 0.048 |
|  | Summer | 0.200 | 0.284 | 0.315 |
| Copper | Monsoon | 1.953 | 0.599 | 0.806 |
|  | Winter | 1.049 | 0.435 | 0.868 |
|  | Summer | 2.175 | 1.455 | 1.716 |
| Lead | Monsoon | 0.304 | 0.145 | 0.186 |
|  | Winter | 0.638 | 0.585 | 0.408 |
|  | Summer | 0.121 | 0.153 | 0.170 |
| Zinc | Monsoon | 0.405 | 0.133 | 0.182 |
|  | Winter | 0.113 | 0.106 | 0.093 |
|  | Summer | 0.608 | 0.808 | 0.513 |
| Nickel | Monsoon | 0.076 | 0.037 | 0.081 |
|  | Winter | 0.029 | 0.034 | 0.045 |
|  | Summer | 0.247 | 0.178 | 0.106 |

**\*All values are expressed in mg/L**

During the course of study it was observed that the values physico-chemical parameters (Alkalinity, Total Hardness, Chlorides, DO, BOD, COD) of the Yamuna River water of Agra city are excessively above the permissible limits prescribed by the recommended agencies WHO (1993), BIS (2003) and ICMR (1975). In the present study, it was observed that the maximum values of alkalinity, Chloride content, total hardness, BOD and COD was found in summer season followed by winter and monsoon season. (Table 3)

The results are similar with the findings of Sikandar (1987), Tiwari (1983) and Tripathi (1982). Lower alkalinity in monsoon season might be due to river water dilution with rain water and higher in summer season due to increase in carbonates and bicarbonate ions. Many researchers like Shinde et al. (2011), Jana (1973), Govindan and Sundaresan (1979), Bheemappa et al. (2015) who reported high value of chlorides in summer and low values during winter and monsoon season. High values in summer may be due to mixing of sewage water, elevation in temperature and evaporation of water (Balasaheb et al. 2015). However, low values in winter may be because of dilution effect and recurrence of water mass changes due to summer stagnation and high rate of sedimentation (Shinde et al. 2011).

Our result are in agreement with the findings of Kataria et al. (1996) who reported maximum value of hardness in summer minimum in winter and moderate in monsoon seasons in Bhopal at Kolar reservoir. Harney et al. (2013) also showed highest values of total hardness during summer which may be attributed to maximum temperatures which enhance the concentration of salts through extreme evaporation.

These results agreed with the findings of Eknath (2013), Mini et al. (2003), Hassan et al. (2017), Tripathi et al. (2015), Kaur and Kaur (2015), Das and Acharya (2003), Maya et al. (2007) and Rao et al. (2013) who reported highest BOD values in summer season and low values in monsoon and winter season. The high values of BOD observed in summer seasons may be due to increase in metabolic rate of numerous microorganism in the degradation of organic waste, low water flow and accumulation of domestic and industrial waste however low values of BOD during monsoon season could be associated with dilution in the water bodies due to huge volume of rain water (Upadhyay and Rana 1991). Our result are in accordance with the findings of Eknath (2013), Mini et al.(2003), Gyananath et al.(2000), Abir (2014), Rao et al. (1990), Harney et al. (2013), Tripathi et al. (2015), Kaur and Kaur (2015) and Hassan et al. (2017) who reported maximum COD in summer and minimum in winter and monsoon season in various water sources. High level of COD values during summer seasons could be attributed to high temperature and increased water evaporation, utilization of oxygen for decomposition of organic matter and discharge of chemical wastes into the Riverine system (Eknath 2013).

During the study period DO was found minimum in summer season. Similar patterns were shown by various researchers; Khanna and Bhutiani (2003) (Ganga River), Singh et al. (2006) (Ganga River) and Badola and Singh (1981) (Alaknanda River). In aquatic systems decreased in DO values during summer is due to increased temperature (Naz and Turkmen 2005). Low values in summer may be due to minimum oxygen holding capacity of water at maximum temperature additionally raised in DO assimilation for biodegradation of organic matter through microbes. However, high values during winter season could be associated with more dissolution of oxygen at minimum temperature of water (Ravindra et al. 2003).

On account of the research work; the Yamuna River water samples contains heavy metal concentrations (Cadmium, Chromium, Copper, Lead and Nickel) above the desirable and admissible levels (WHO, 2008 and BIS 2012) except for Zinc. Throughout the study, highest concentrations of heavy metals (Cadmium, Chromium, Copper, Zinc and Nickel) were present in summer season and lowest concentration occurred in winter and monsoon season. This was supported by Zyadah (1995) who reported that the seasonal fluctuations in heavy metals concentration might be due to variation in the quantity of agricultural drainage water, industrial and sewage effluents discharged into the aquatic systems. Similarly, Abdel-Moati and El-Sammak (1997) and Olatunji and Osibanjo (2012) explained that during the study period, the maximum concentration of heavy metals in the water samples during summer season is because of decrease in water levels in Rivers which results in the increase concentration of the heavy metals. Researchers Sophia et al. (2017), Ibrahim and Omar (2013), Shanbehzadeh et al. (2014), Kar et al. (2008) and TEKIN-OZAN (2015) also revealed that the heavy metals contamination in the water was found maximum during summer or dry season.

### Non-Culturable Bacteria

To unravel the non-culturable bacterial diversity in Yamuna river water collected from different sites (Site-1 Kailash Ghat, Site-2 Poiyah Ghat and Site-3 Hathi Ghat) of Agra, uniform suspension of all three water samples were prepared by mixing them thoroughly for the purpose of metagenomic analysis (at-Xcelris Labs Limited, Ahmedabad). The DNA was isolated from the samples and run on 0.8% gel for further metagenomic based analysis. The DNA obtained was quantified using Qubit Fluorometer and the result showed a good quality of DNA in the samples with 3.30 μg yield and 41.2 ng/μl concentration.

### Whole Metagenomic Analysis: Data Statistics

The libraries were prepared from the water sample by *Truseq Nano DNA Library preparation kit*. The average sizes of libraries were 464bp. The library was sequenced on Illumina platform (2 × 150 bp chemistry) to generate ∼3 GB data/Sample. Collectively, there were about 23409644 reads obtained from the sample. *De novo* assembly of high quality PE reads was accomplished using CLC Genomics Workbench 6.0 at default parameters (Deletion cost: 3, Automatic word size: Yes, Perform scaffolding: Yes, Insertion cost: 3 Minimum contig length: 200, Length fraction: 0.5, Mismatch cost: 2, Similarity fraction: 0.8). The statistical elements of the assemblies were calculated by in house pearl scripts and the results are given in the Table 4.

**Table 4:** Assembly Statistics.

| Assembly Element | Water sample |
| --- | --- |
| Number of scaffolds | 144121 |
| Total genome length in Mb | 135.45 |
| average scaffold size | 940 |
| N50 | 884 |
| Max scaffold size | 209613 |

### Gene Prediction

High quality reads were further used for denovo assembly with CLC. The assembly resulted in 144121 scaffolds for Water sample. Total of 240310 genes were predicted from the scaffolds of Water sample using Prodigal (v2.6.3). These genes predicted were considered for downstream analysis. From the Water sample genes and the predicted genes from scaffolds were then taken for taxonomic and functional analysis using Kaiju, Cognizer respectively (Table 5).

**Table 5:**
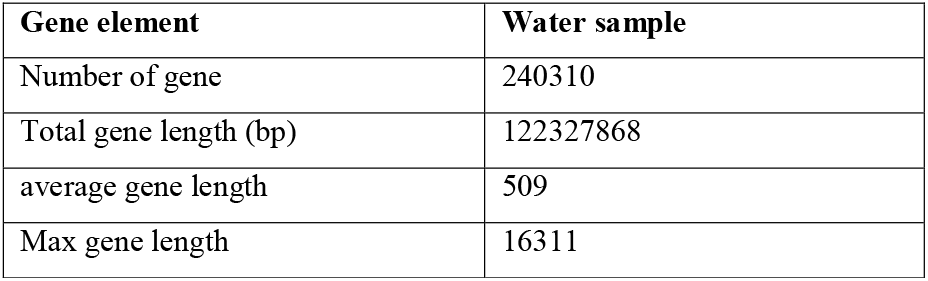
Statistics of predicted ORFs.

| Gene element | Water sample |
| --- | --- |
| Number of gene | 240310 |
| Total gene length (bp) | 122327868 |
| average gene length | 509 |
| Max gene length | 16311 |

### Taxonomic Annotation using Kaiju

Kaiju is a fast, free and sensitive metagenome classifier which finds maximum exact matches on the protein-level using the Burrows–Wheeler transform algorithm. Kaiju classified individual metagenomic reads through a reference database consisting the annotated protein-coding genes of a set of microbial genomes. By default, Kaiju uses either the microbial subset of the non-redundant protein database nr used by NCBI BLAST or the available complete genomes from NCBI RefSeq, optionally also including fungi and microbial eukaryotes. The final sequences were uploaded to the Kaiju webserver (http://kaiju.binf.ku.dk) with the parameters as run mode (greedy), minimum match length (11), maximum match score (65), allowed mismatches (5), SEQ low complexity filter (yes), etc.

### Taxonomic distribution of Water sample at different levels

The taxonomic classification of bacterial community and their abundance in the sample was done with a similarity search of 23409644 reads. Bubble size indicates taxon abundance relative to its maximum abundance (largest bubble size). The size of the circle is scaled logarithmically to represent the number of sequences assigned directly to the taxon. From the Fig. 2 it can be inferred that Proteobacteria was found to be the most abundant. The sample of presented study possessed a dominating population of Proteobacteria species followed by Bacteroidetes, Verrucomicrobia, Actinobacteria, and so on.

**Fig. 2.**
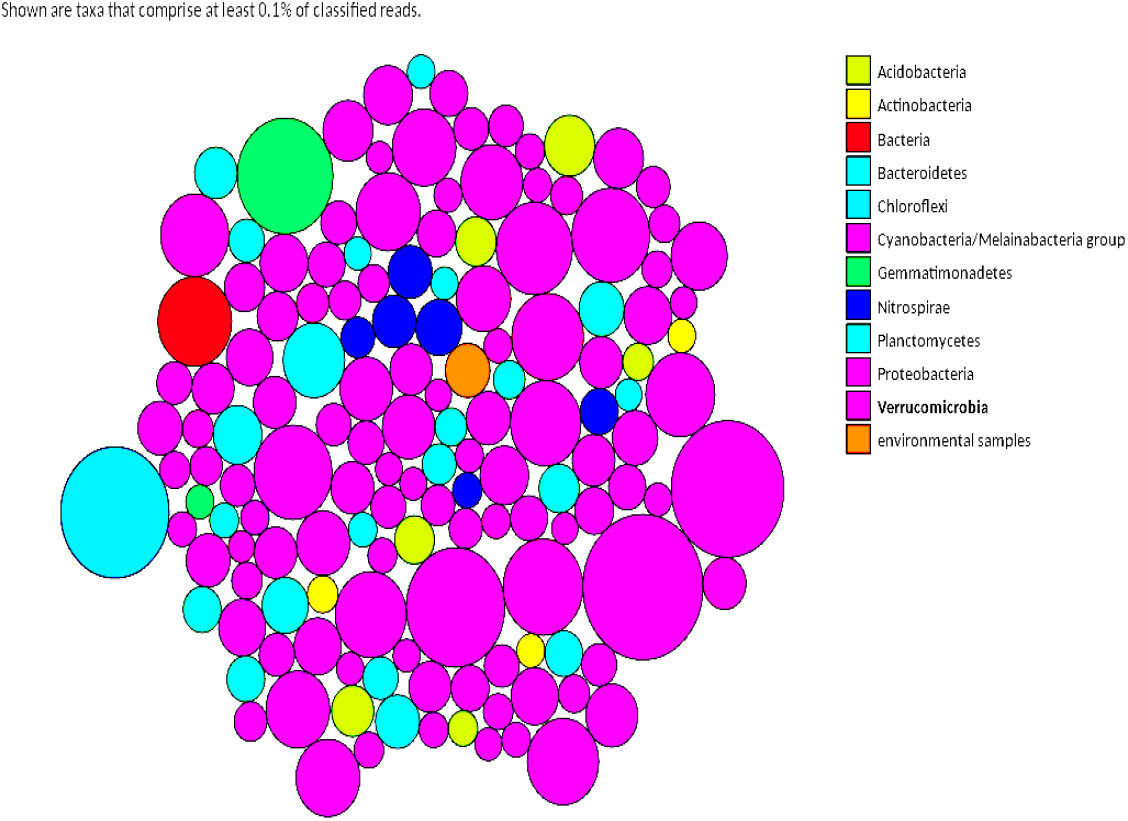
Bubble plot showing the relative taxonomic abundance of Water sample.

Taxonomic hits distribution of the top 50 phylum, class, order, family, genus and species shows that Water sample has 85850 Proteobacteria followed by 13129 Bacteroidetes (Fig. 3). Proteobacteria and Bacteriodetes are often the most abundant phylum found in the aquatic systems (Eilers et al. 2000). Remaining classified phyla with > 1000 relative abundance were found in Verrucomicrobia (7365), Actinobacteria (6525), Planctomycetes (6051), Chloroflexi (4250), Acidobacteria (3608), Gemmatimonadetes (2900), Nitrospirae (2605), Firmicutes (2327) in the water sample. Other remaining phyla from *Candidatus Omnitrophica* to *Candidatus Levybacteria* were < 1000 in abundance. At the class level, taxonomic hit distribution of top 50 classes showed that water sample has 37030 Betaproteobacteria followed by Gammaproteobacteria (20632), Alphaproteobacteria (20435), Deltaproteobacteria (6293), Actinobacteria (5015) and Planctomycetia (5389). Thirteen classes were found with abundance between (500-5000). The remaining classes have abundance less than 500 as given in Fig. 4.

**Fig. 3.**
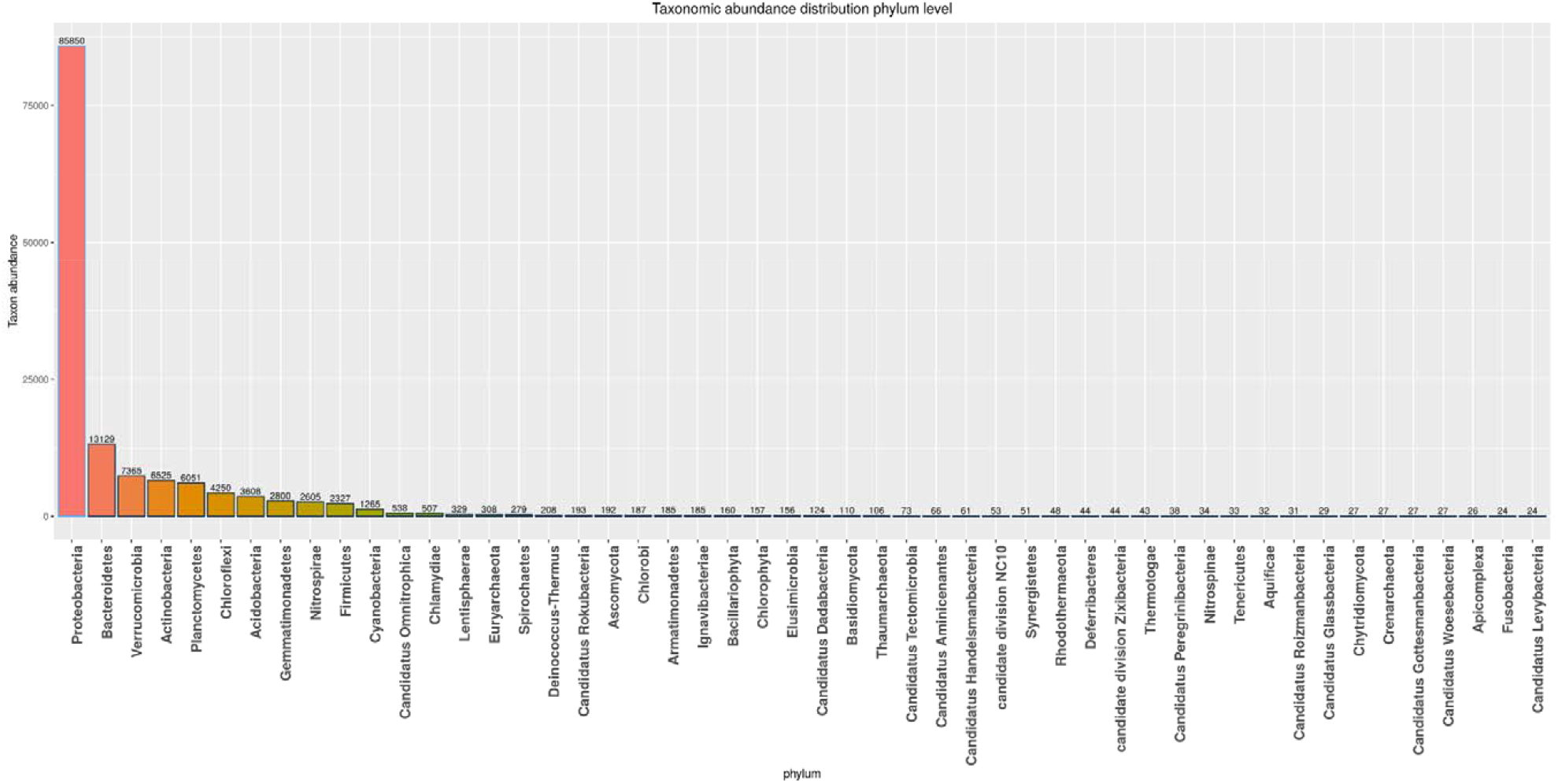
Bar chart showing the taxonomic abundance of Water sample at phylum level. From the figure it can be inferred that Proteobacteria is the most abundant phylum followed by Bacteroidetes

**Fig. 4.**
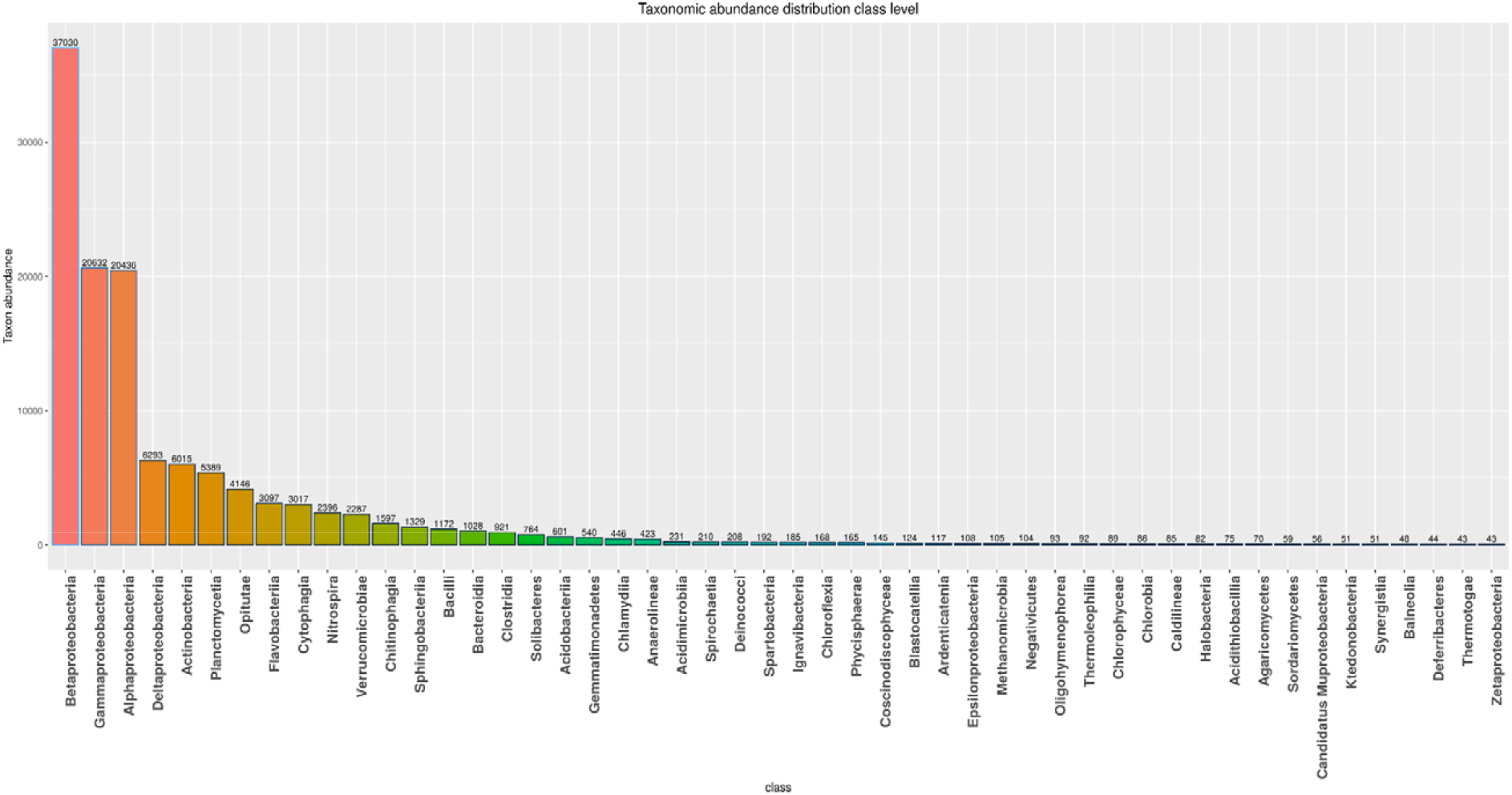
Bar chart showing the taxonomic abundance of Water sample at class level. From the figure it can be inferred that Betaproteobacteria is the most abundant class followed by Gammaproteobacteria

At order level, the 50 taxonomic hits distribution was 20033 Burkholderiales followed by 9505 Rhodocyclales (Fig. 5), 11070 Comamonadaceae followed by 9408 Rhodocyclaceae (Fig. 6) for family, 4853 Hydrogenophaga followed by 4684 Thauera (Fig. 7) for genus level. It is found that various generas related to Proteobacteria and family Comamonadaceae are involved in degradation of polycyclic aromatic hydrocarbons and iron oxidation (Emerson et al. 2015; Wang et al. 2016). The abundance of bacterial community in water samples shows the presence of the pollutants (PAH and others). Fig. 7 shows the taxonomic classification depicting top 50 genera for the tested sample. At genera level out of all metagenomic reads 4853 were Hydrogenophaga followed by 4684 Thauera. Also, Nitrosomonas, Solimonas, Pseudomonas, Magnetospirillum, Nitrospira, Azoarcus, Optitutus, etc. were some of the most dominating genera in the sample.

**Fig. 5.**
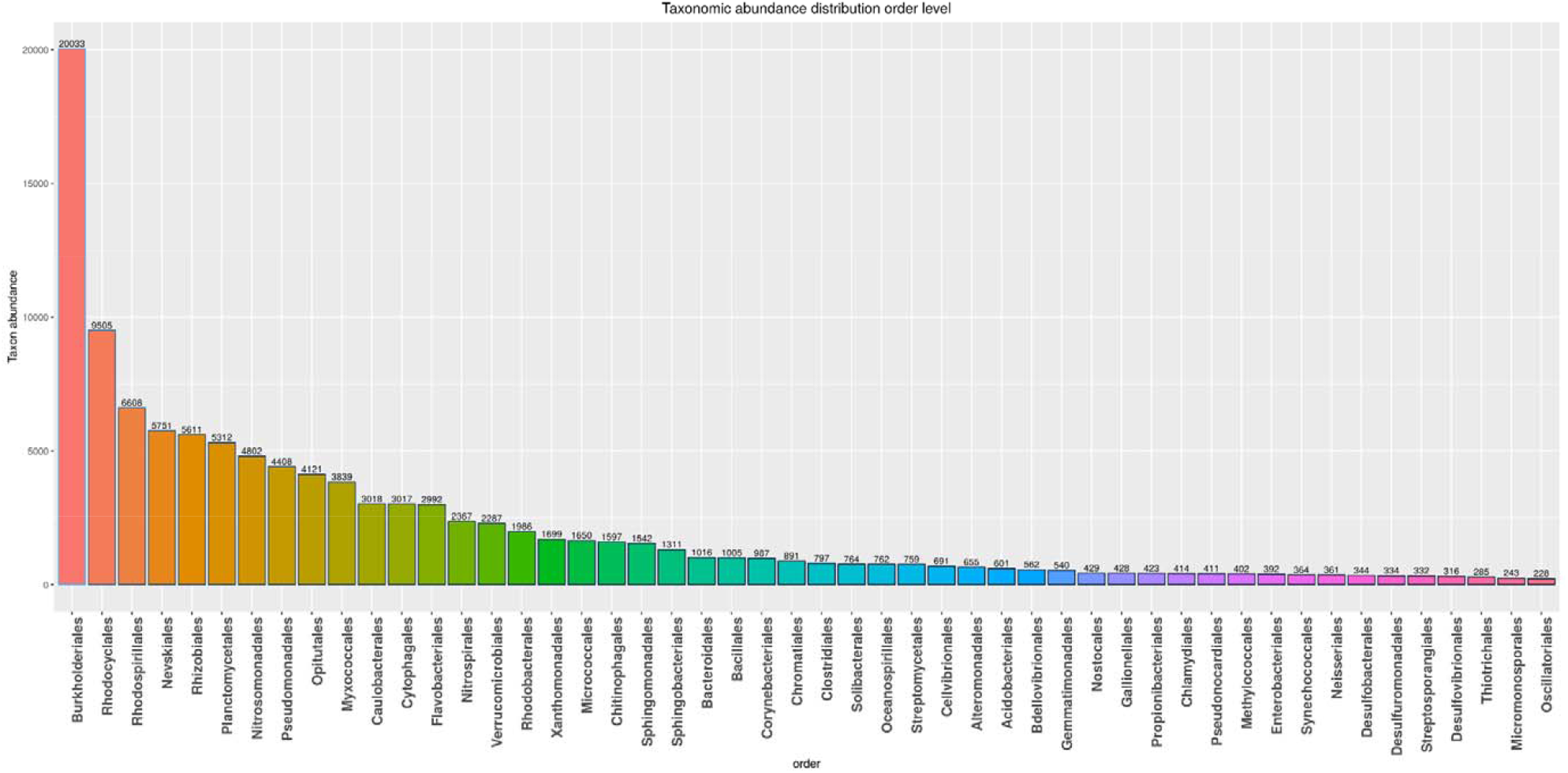
Bar chart showing the taxonomic abundance of Water sample at order level. From the figure it can be inferred that Burkholderiales is the most abundant order followed by Rhodocyclales.

**Fig. 6.**
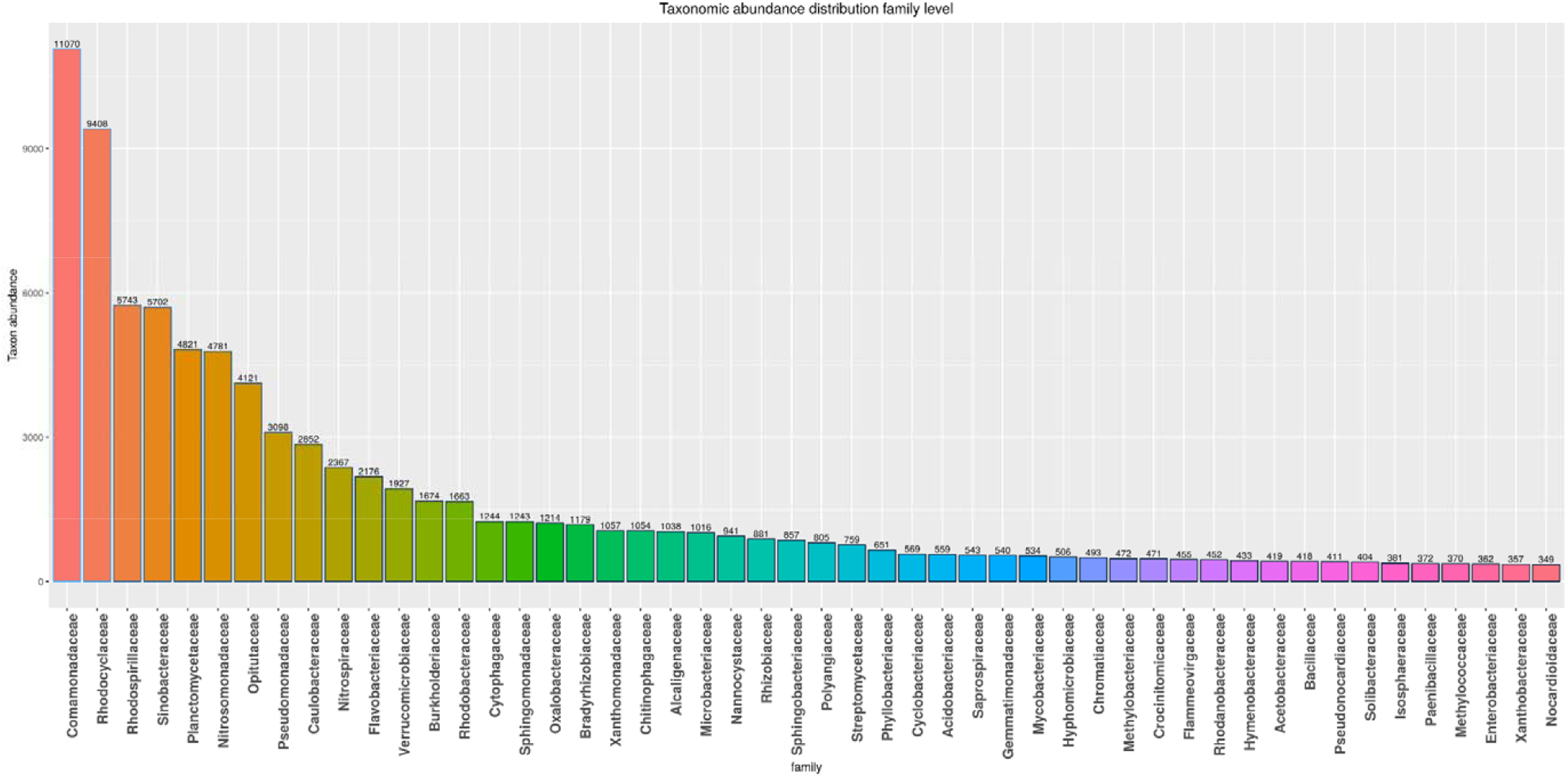
Bar chart showing the taxonomic abundance of Water sample at family level. From the figure it can be inferred that Comamonadaceae is the most abundant family followed by Rhodocyclaceae.

**Fig. 7.**
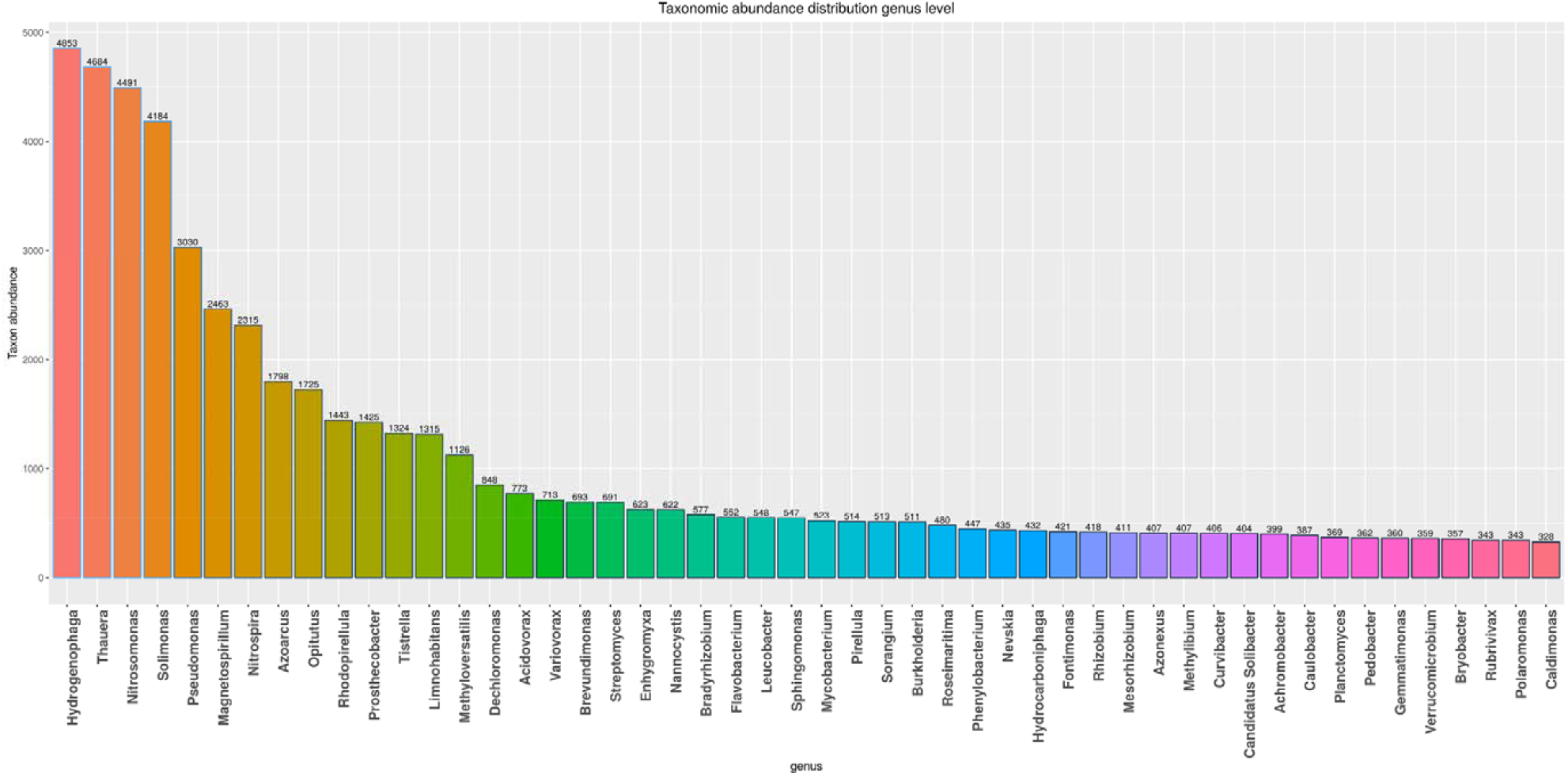
Bar chart showing the taxonomic abundance of Water sample at genus level. From the figure it can be inferred that *Hydrogenophaga* is the most abundant genus followed by *Thauera*.

At species level, 2630 *Chloroflexi bacterium OLB14* was followed by 2050 *Gemmatimonadetes bacterium SCN 70* and upto 318 (*Plesiocystis pacifica*). No species were found <200 abundance in the distribution (Fig. 8). This *Chloriflexi bacterium* is seen responsible for dechlorination of polychlorinated biphenyls (PCB) (Zanaroli et al. 2012). Taxonomic analysis results can be summarized in the following order, Proteobacteria, Betaproteobacteria, Burkholderiales, Comamonadaceae, Hydrogenophaga, *Chloroflexi bacterium OLB14*.

**Fig. 8.**
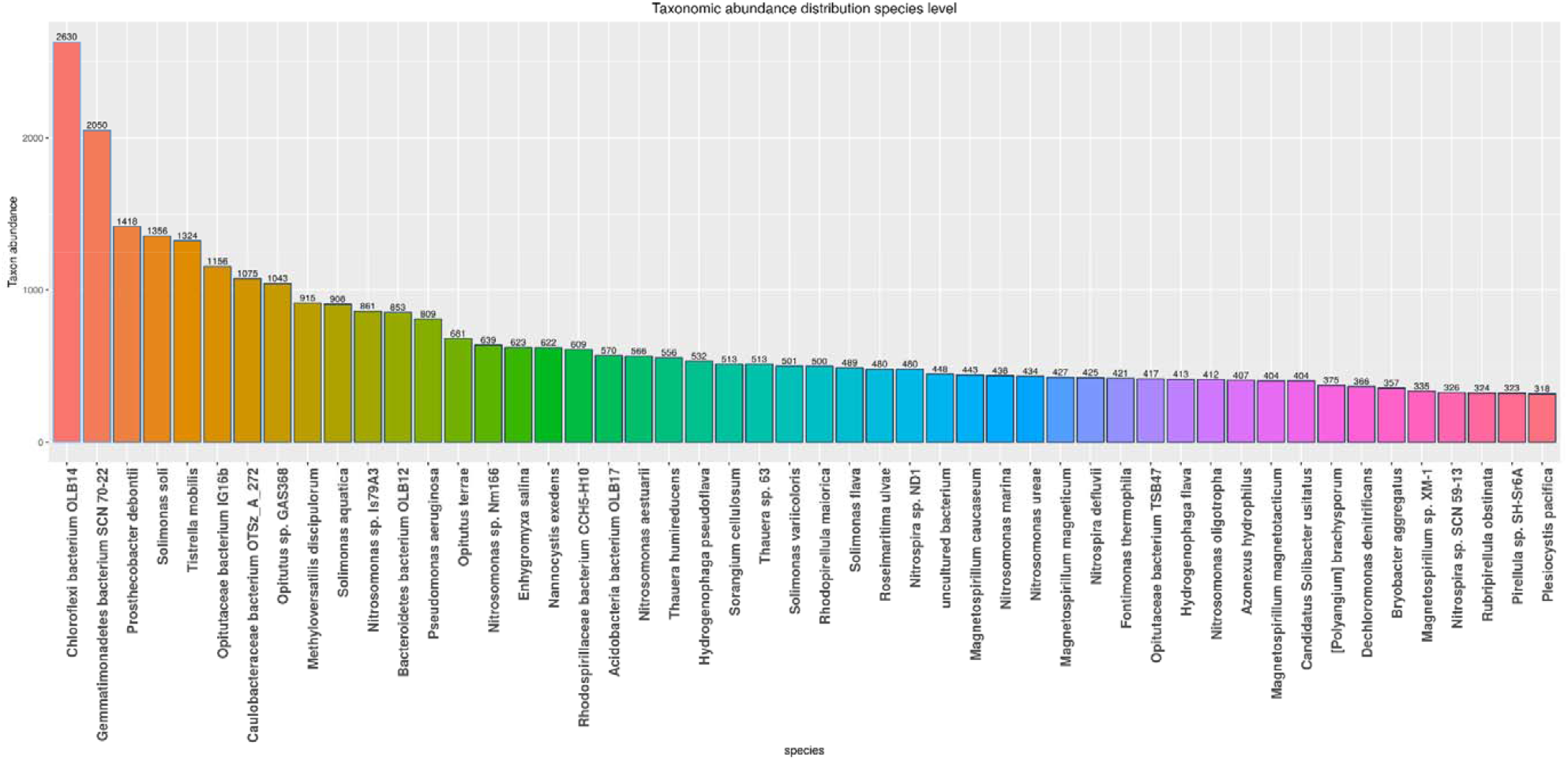
Bar chart showing the taxonomic abundance of Water sample at species level. From the figure it can be inferred that *chloroflexi bacterium OLB14* is the most abundant genus followed by *gemmatimonadetes bacterium SCN 70*.

### Functional Annotation using Cognizer

COGNIZER was used to assess the functional capacities of microbial communities present in the samples. Cognizer is a comprehensive stand-alone framework which is enabled to simultaneously provide COG, KEGG (Kyoto Encyclopedia of Genes and Genomes), Pfam, GO and FIGfams annotations to individual sequences constituting metagenomic datasets. In total 240310 predicted genes containing scaffolds of the water sample were considered for the final downstream analysis using Cognizer. Our metagenome sequence has been uploaded on the NCBI website under the accession number PRJNA608246. Fig. 9 shows the number of sequences in water sample dataset that were annotated by the different databases. These include protein databases, protein databases with functional hierarchy information etc. The maximum number of hits was observed against GO, followed by KEGG, Pfam, COG and FIG database (Table 6).

**Table 6:** Functional category classifications results can be summarized as mentioned below.

| Description | Water sample |
| --- | --- |
| GO | 160247 |
| KEGG | 140006 |
| Pfam | 125840 |
| COG | 116675 |
| FIG | 76857 |

**Fig. 9.**
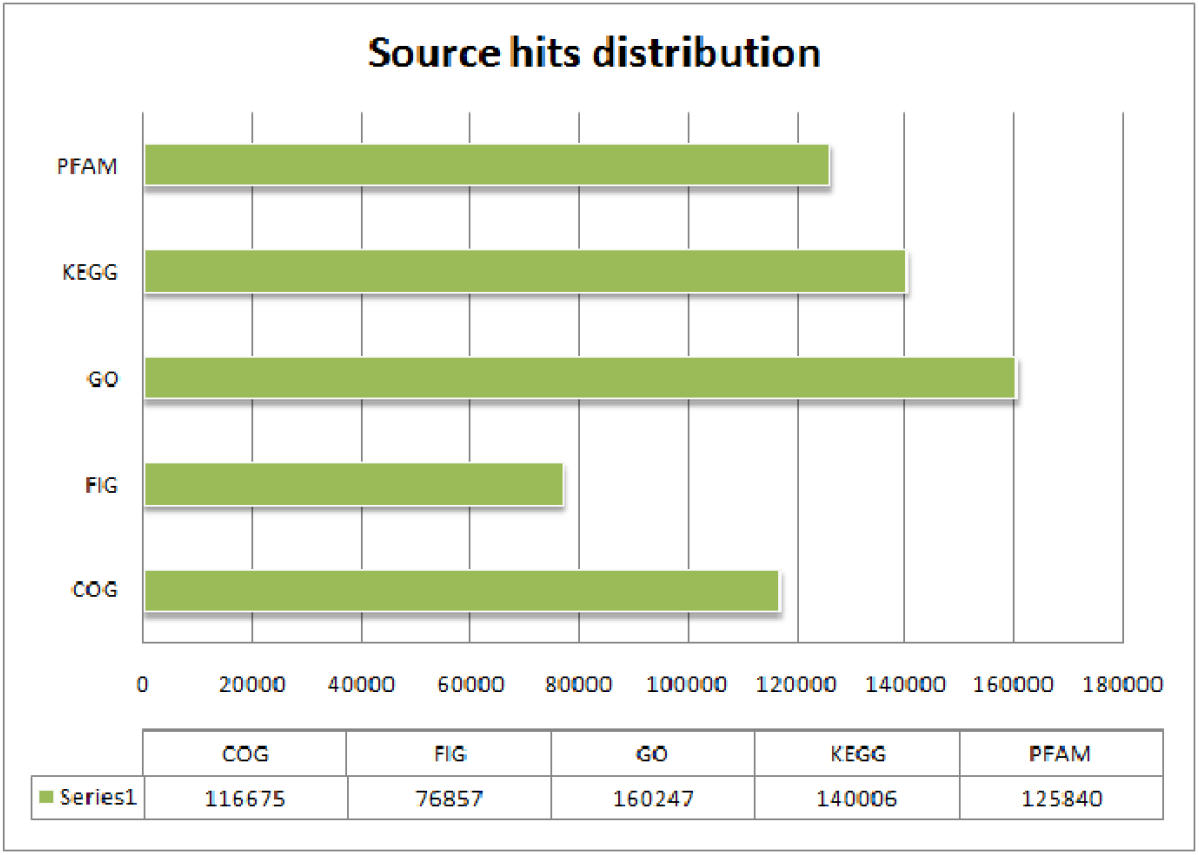
Source hits distribution of Water sample. Each number indicated represents the number of sequences annotated against the respective databases. Maximum number of hits was observed in GO database, followed by KEGG and PFAM.

GO functional analysis has 160247 predicted functions against GO database.

### COG Functional Category Hits Distribution

COG (Clusters of Orthologous Genes) database also known as Clusters of orthologous groups of proteins which contain gene products related to a wide range of bacteria and archaea species. This database include gene related to various signaling enzymes, translational factors, RNA processing and transduction. COG functional Category has 116675 predicted functions, out of which 13569 are for general function prediction only and 10428 are for Amino acid transport and metabolism as shown in the Fig. 10. Similarly, other predicted genes for function such as defense mechanism, signal transduction, cell wall/membrane/envelope biogenesis, replication recombination and repair, etc. were found in the results.

**Fig. 10.**
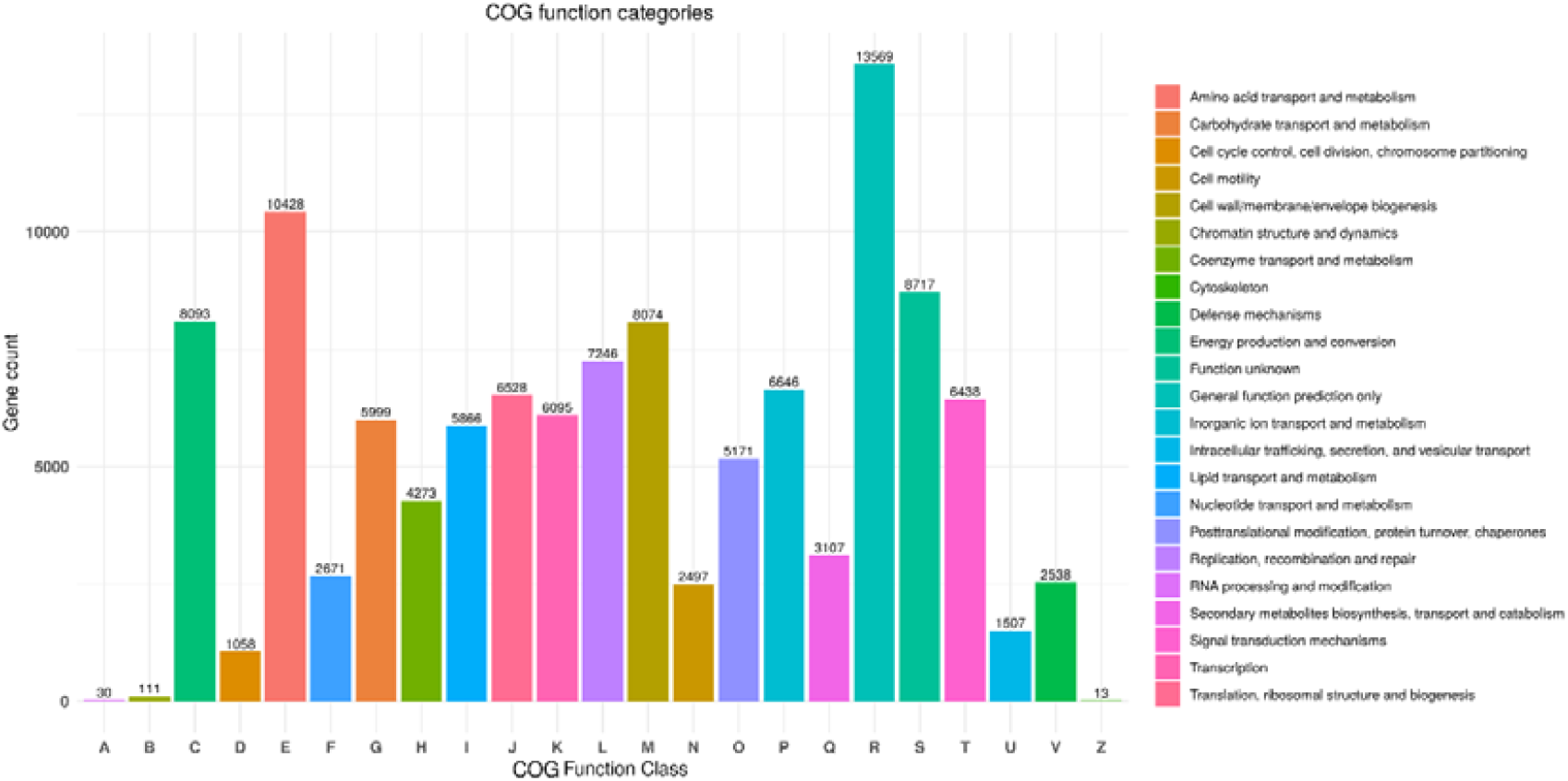
COG Functional Category Hits Distribution of Water sample

### KO Functional Category Hits Distribution

KEGG database interprets the genomic data to its functional forms and other high throughput information. The protein and gene functions are related to the ortholog groups (OG) which are stored in the KEGG Orthology (KO) database. The KEGG database has complete genome sequences that help to link set of genes with functions of the organism. (Kanehisa et al. 2016). In this study, KO Functional Category has 140006 predicted functions in which most of the sequence falls in Metabolism followed by Environmental information processing function as shown in the Fig.11.

**Fig. 11.**
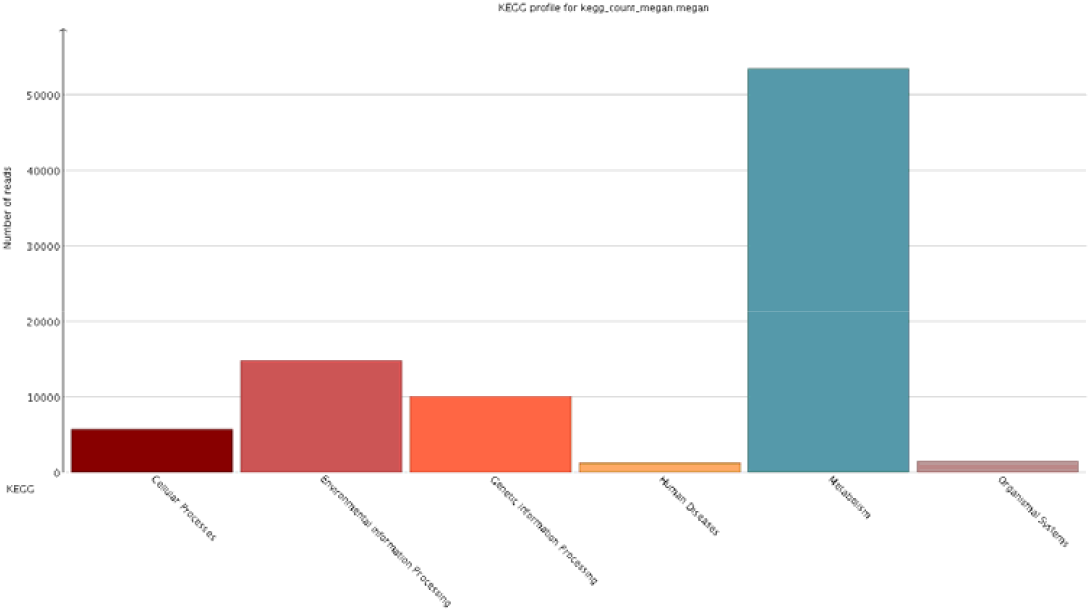
KO Functional Category Hits Distribution of Water sample

## DISCUSSION AND CONCLUSION

Rivers have played a very important role in development of civilization, culture, settlement of urban area thus it plays a critical and crucial role in the prosperity of a nation affecting the different aspects of its economic status. In India’s religious literature and ancient scriptures great respect and spirituality has been expressed for Yamuna river but now-a days the holy river has fallen sick due to civic indifference and poor governance. River Yamuna is one of the most sacred still most polluted rivers of India.

There are many metagenomic researches done on polluted water sample to identify the bacterial contamination (Ding et al. 2017; Saleem et al. 2018; Samson et al. 2019; Behera et al. 2020). This study of metagenomic sequencing of water sample from Yamuna River revealed different organisms for the evaluation of microbial diversity and abundance. The results showed that Proteobacteria (phylum), Betaproteobacteria (class), Burkholderiales (order), Comamonadaceae (family), Hydrogenophaga (genus) and *Chloroflexi bacterium* OLB 14 (species) being the most dominant bacterial taxonomic abundance in the Yamuna river water samples. The major phylum (Proteobacteria) with predominant class of Alpha-, Beta-, Gamma-, and Delta-Proteobacteria were present in the sample in high abundance. This is similar to other studies performed on freshwater systems (Tsagaraki et al. 2018). The second most abundant class was of Gamma-Proteobacteria, these are facultative anaerobes which contribute in nitrogen cycling and vast metabolic diversity. Delta-Proteobacteria was the fourth abundant class found which consist of members capable of reducing iron, sulphate and dehalogentaion (Sandford et al. 2002). The dominance of these classes in water indicates towards the large scale domestic, industrial and human activities that have provided favorable environment for such anaerobes to grow.

A similar study has been done on Yamuna River at Kalindi Kunj, Delhi in which Proteobacteria (33.68%) and Bacteriodetes (24.07%) showed maximum abundance at phylum level. Similar results were also observed by Wang *et al*. (2014), Miyashita (2015) and Liu *et al*. (2012) who reported Proteobacteria as the most prevalent phylum found in domestic sewage water, drinking water as well as in soil. Many researchers (Kent et al. 2001; Li et al. 2006 and Schnetzer et al. 2011) reported the dominance of Proteobacteria in wastewaters and soils. α-Proteobacteria is the dominant Proteobacterial class in the marine microbial community found at the surface of the Yellow Sea (Bai et al. 2009; Sogin et al. 2006).

Metagenomic study for identification of microbial communities in water helps in the exploration of novel genes with different metabolic functions related to respective organism. The alarming levels of pollution in rivers necessitate the functional analysis of microbial communities. In the present study, the functional annotation of genes have shown maximum hit with GO databases, followed by KEGG or KO and PFAM. 140006 sequences were annoted with KO database. The functional orthologies found with KO indicates the bacterial community capable of degrading organic and inorganic pollutants. These results could conclude that sample may be rich in hazardous pollutants. The analysis of KEGG related genes revealed their importance in metabolic functions, processing of environmental and genetic information, cellular processes, etc. Also, some gene sequences were found related to the human infection pathways. PFAM functional analysis has 125840 predicted functions against a manually curated Pfam database. Pfam functional annotation is used to assess protein families and domain. Also, FIGfams functional analysis has 76857 predicted functions against manually annotated FIGfams database. FIGfams functional annotation is used to find the families of proteins that have the same functions. Hence, we can that a substantial bacterial diversity was observed in the Yamuna river water samples collected from different sites.

From the results and observation it can be concluded that the quality of Yamuna river water of Agra city is highly contaminated at all sampling sites that may be due to small and large scale industries, disposal of untreated sewage and agricultural sectors. This study is helpful in addressing the issues of polluting rivers contaminated with pathogenic bacteria and the respective pollutants. The information will help in making of effective plans in order to improve the water quality for its consumption. Also, the study helps in understanding of different microbial communities or microbial population dynamics in the contaminated water which cannot be cultured and help in the development of future remediation strategies related to them.

## ACKNOWLEDGEMENT

The authors would like to acknowledge the support of Xcelris Labs for extending support, wherever required during the research and UGC SAP for financial help during the research studies.

## ETHICAL STATEMENT

I testify on behalf of all co-authors that our research article “Metagenomics Study of Bacterial Communities from Yamuna River water of city of Taj, Agra” has not been published elsewhere.

